# SQUID: Transcriptomic Structural Variation Detection from RNA-seq

**DOI:** 10.1101/162776

**Authors:** Cong Ma, Mingfu Shao, Carl Kingsford

## Abstract

Transcripts are frequently modified by structural variations, which leads to a fused transcript of either multiple genes (known as a fusion gene) or a gene and a previously non-transcribing sequence. Detecting these modifications (called transcriptomic structural variations, or TSVs), especially in cancer tumor sequencing, is an important and challenging computational problem. We introduce SQUID, a novel algorithm to accurately predict both fusion-gene and non-fusion-gene TSVs from RNA-seq alignments. SQUID unifies both concordant and discordant read alignments into one model, and doubles the accuracy on simulation data compared to other approaches. With SQUID, we identified novel non-fusion-gene TSVs on TCGA samples.

## 1 Background

Large-scale transcriptome sequence changes are known to be associated with cancer [1, 2]. Those changes are usually a consequence of genomic structural variation (SV). By pulling different genomic regions together or separating one region into pieces, structural variants can potentially cause severe alteration to transcribed or translated products. Transcriptome changes induced by genomic SVs, called transcriptomic structural variants (TSVs), can have a particularly large impact on disease genesis and progression. In some cases, TSVs bring regions from one gene next to regions of another, causing exons from both genes to be transcribed into a single transcript (known as a fusion gene). Domains of the corresponding RNA or proteins can be fused, inducing new functions or causing loss of function, or the transcription or translation levels can be altered, leading to disease states. For example, *BCR-ABL1* is a well-known fusion oncogene for chronic myeloid leukemia [3], and the *TMPRSS2-ERG* fusion product leads to over-expression of *ERG* and helps triggers prostate cancer [4]. These fusion events are used as biomakers for early diagnosis or treatment targets [5]. In other cases, TSVs can affect genes by causing a previously non-transcribed region to be incorporated into a gene, causing disruption to the function of the altered gene. There are fewer studies on these TSVs between transcribed and non-transcribed regions, but their ability to alter downstream RNA and protein structure is likely to lead to similar results as fusion gene TSVs.

Genomic SVs are typically detected from whole-genome sequencing (WGS) data by identifying reads and read pairs that are incompatible with a reference genome [e.g., 6–11]. However, WGS data are not completely suitable to infer TSVs since they neither inform which region is transcribed nor reveal how the transcribed sequence will change if SVs alter a splicing site or the stop codon. In addition, WGS data is more scarce and more expensive to obtain than RNA-seq [12] measurements, which target transcribed regions directly. RNA-seq is relatively inexpensive, high-throughput, and widely available in many existing and growing data repositories. For example, The Cancer Genome Atlas (TCGA, https://cancergenome.nih.gov) contains RNA-seq measurements from thousands of tumor sample across various cancer types, but 80% of tumor samples in TCGA have RNA-seq data but no WGS data (Supplementary Figure S1). While methods exist to detect fusion genes from RNA-seq measurements [e.g., 13–21], fusion genes are only a subset of TSVs, and existing fusion gene detection methods rely heavily on current gene annotations and are generally not able or at least not optimized to predict non-fusion-gene TSV events. The idea of de novo transcript assembly [e.g., 22–25] followed by transcript-to-genome alignment [e.g., 26–28] is used in some fusion-gene detection methods. These approaches rely on annotation-based filtering steps to achieve the high accuracy. Although it is possible to extend these approaches to non-fusion-gene TSV detection, the lack of annotation information for non-transcribing regions makes these approaches less suitable for finding non-fusion-gene TSV. This motivates the need for a method to detect both types of TSVs directly from RNA-seq data.

We present SQUID, the first computational tool that is designed to comprehensively predict both types of TSVs from RNA-seq data. SQUID divides the reference genome into segments and builds a genome segment graph from both concordant and discordant RNA-seq read alignments. In this way, it can detect both fusion-gene events and TSVs incorporating previously non-transcribed regions into transcripts. Using an efficient, novel integer linear program (ILP), SQUID rearranges the segments of the reference genome so that as many read alignments as possible are concordant with the rearranged sequence. TSVs are represented by pairs of breakpoints realized by the rearrangement. Discordant reads that cannot be made concordant through the optimal rearrangement given by the ILP are discarded as false positive discordant reads, likely due to misalignments. By building a consistent model of the entire rearranged genome and maximizing the number of overall concordant read alignments, SQUID drastically reduces the number of spurious TSVs reported compared with other methods.

SQUID features high accuracy. SQUID is usually > 20% more accurate than applying WGS-based SV detection methods to RNA-seq data directly. It is similarly more accurate than the pipeline that uses de novo transcript assembly and transcript-to-genome alignment to detect TSVs. We also show that SQUID is able to detect more TSVs involving non-transcribed regions than any existing fusion gene detection method.

We use SQUID to detect TSVs within 401 TCGA tumor samples of four cancer types (99-101 samples each of breast invasive carcinoma [29], bladder urothelial carcinoma [30], lung adenocarcinoma [31], and prostate adenocarcinoma [32]). SQUID’s predictions suggest that breast invasive carcinoma has more samples with a larger or smaller number of TSVs / non-fusion-gene TSVs than other cancer types. We also characterize the differences between fusion-gene TSVs and non-fusion-gene TSVs. Non-fusion-gene TSVs, for example, are more likely to be intra-chromosomal events. We show that breakpoints can occur in multiple samples, and among those that do repeatedly occur, their breakpoint partners are also often conserved. Finally, we identify several novel non-fusion-gene TSVs that affect known tumor suppressor genes, which may result in loss-of-function of corresponding proteins and play a role in tumor genesis.

## 2 Results

### 2.1 A novel algorithm for detecting TSVs from RNA-seq

SQUID predicts TSVs from RNA-seq alignments to the genome (Figure 1 provides an overview). To do this, it seeks to rearrange the reference genome to make as many of the observed alignments consistent with the rearranged genome as possible. Formally, SQUID constructs a graph from the alignments where the nodes represent boundaries of genome segments and the edges represent adjacencies implied by the alignments. These edges represent both concordant and discordant alignments, where concordant alignments are those consistent with the reference genome and discordant alignments are those that are not. SQUID then uses a novel integer linear program (Section 4.2) to order and orient the vertices of the graph to make as many edges consistent as possible. Adjacencies that are present in this rearranged genome but not present in the original reference are proposed as predicted TSVs. The identification of concordant and discordant alignments (Section 4.3), construction of the genome segments (Section 4.4), creation of the graph, and the reordering objective function (Section 4.1) are described in the Methods section.

**Figure 1:**
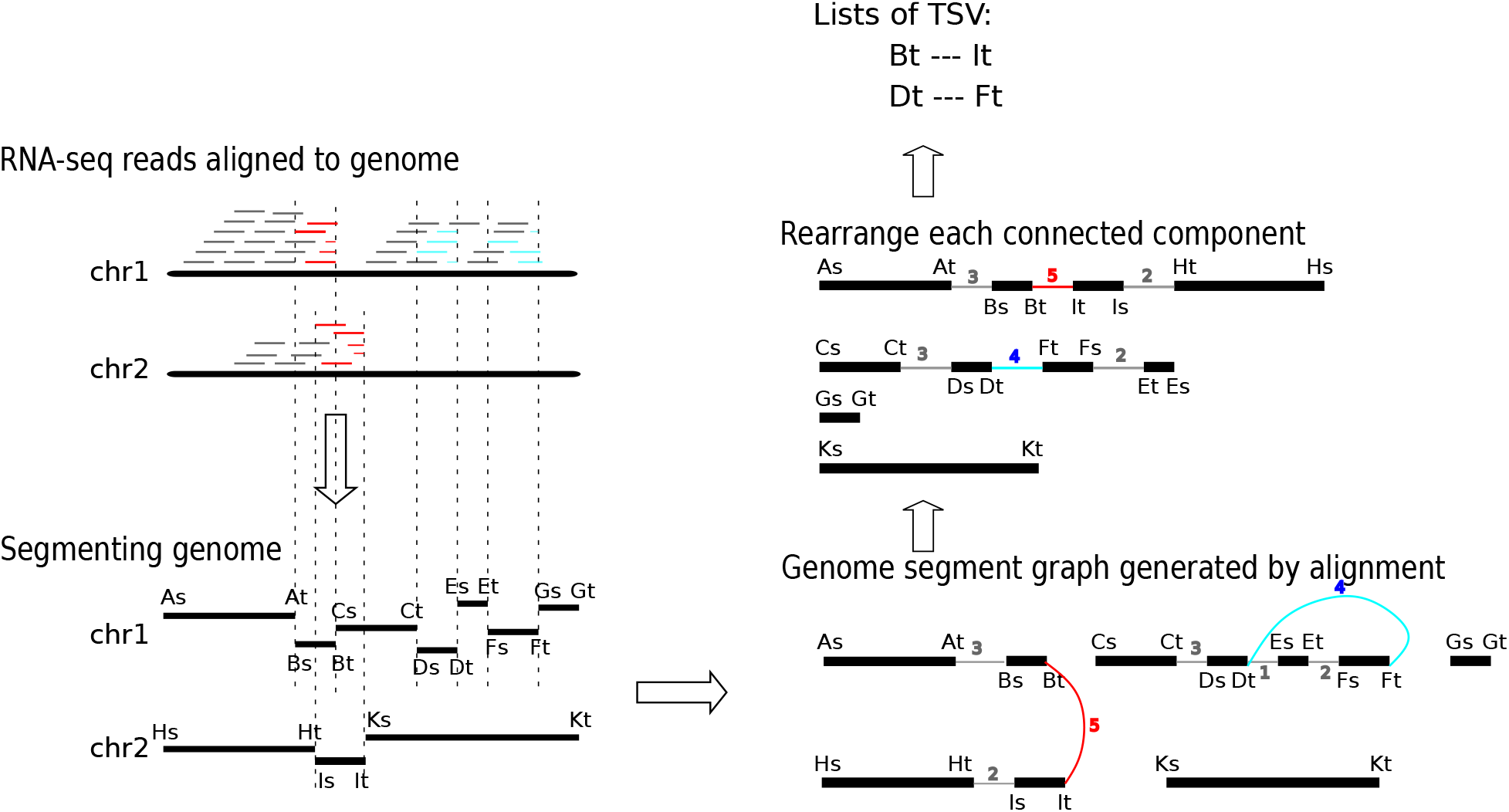
Overview of the SQUID algorithm. Based on the alignments of RNA-seq reads to the reference genome, SQUID partitions the genome into segments, connects the endpoints of the segments to indicate the actual adjacency in transcript, and finally reorders the endpoints along the most reliable path. Each edge in the final path that comes from discordant read alignments represents a TSV.

### 2.2 SQUID is accurate on simulation data

Overall, SQUID’s predictions of TSVs are far more precise than other approaches at similar sensitivity on simulated data (Section 4.7). SQUID achieves 60% to 80% percent precision and about 50% percent sensitivity on simulation data (Figure 2). SQUID’s precision is > 20% higher than several de novo transcriptome assembly and transcript-to-genome alignment pipelines (for details see Supplementary Text), and the precision of WGS-based SV detection methods on RNA-seq data is even lower. The sensitivity of SQUID is similar to de novo assembly with MUMmer3 [26], but a little lower than DELLY2 [6] and LUMPY [7] with SpeedSeq [33] aligner. The overall sensitivity is not as high as precision, which is probably because there are not enough supporting reads aligned correctly to some TSV breakpoints. The fact that assembly and WGS-based SV detection methods achieve similar sensitivity corroborates the hypothesis that it is the data limiting the achievable sensitivity.

**Figure 2:**
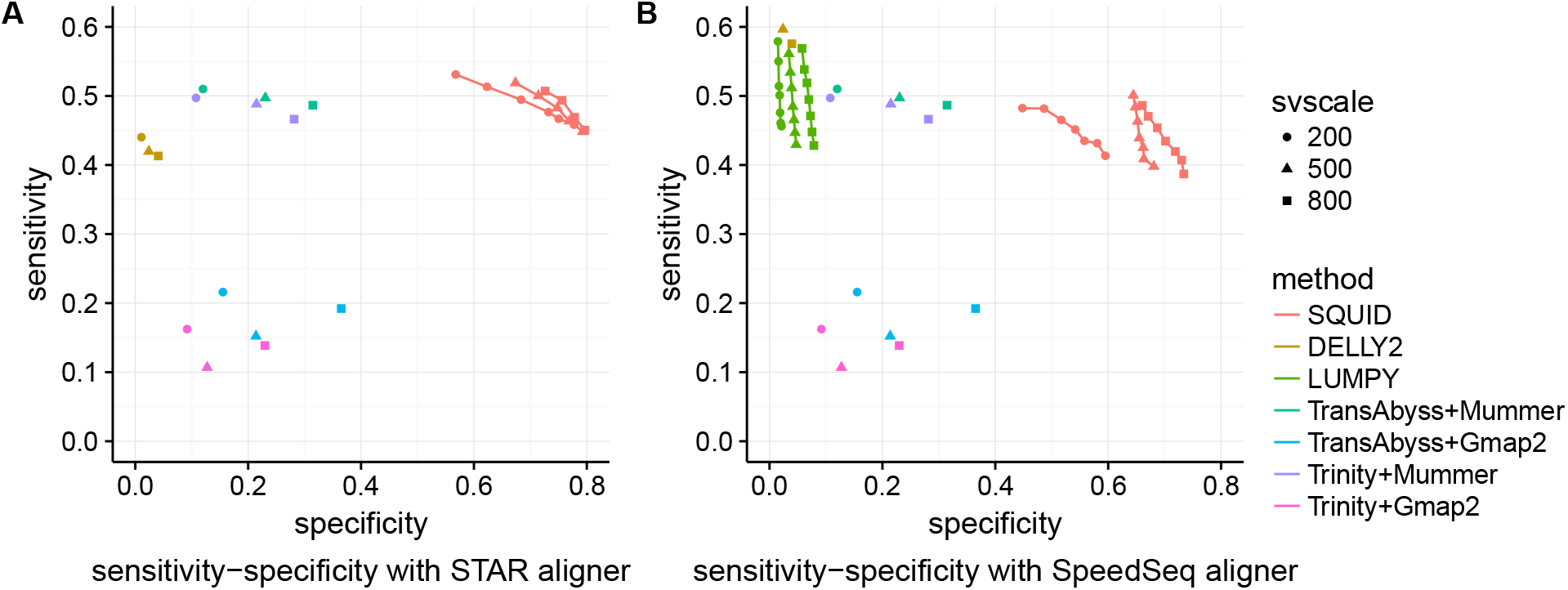
Performance of SQUID and other methods on simulation data. Different number of SVs (200, 500, 800 SVs) are simulated in each dataset. Each simulated read is aligned with both (A) STAR and (B) SpeedSeq aligner. If the method allows for user-defined minimum read support for prediction, we vary the threshold from 3 to 9, and plot a curve on sensitivity-specificity curve (SQUID and LUMPY), otherwise it is shown as a single point

The low specificity of the pipeline- and WGS-based methods shows neither of these types of approaches are suitable for TSV detection from RNA-seq data. WGS-based SV detection methods are able to detect TSV signals, but not able to filter out false positives. Assembly-based approaches require solving the transcriptome assembly problem which is a harder and more time-consuming problem, and thus errors are more easily introduced. Further, the performance of assembly pipelines depends heavily on the choice of software — for example, MUMmer3 [26] is better at discordantly aligning transcripts than GMAP [27]. Dissect [28] is another transcript-to-genome alignment method that is designed for the case where SVs exist. (Unfortunately, Dissect did not run to completion on the some of the dataset tested here.) It is possible that different combinations of de novo transcript assembly and transcript-to-genome alignment tools can improve the accuracy of the pipelines, but optimizing the pipeline is out of scope of this work.

SQUID’s effectiveness is likely due to its unified model of both concordant reads and discordant reads. Coverage in RNA-seq alignment is proportional to the expression level of the transcript, and using one read count threshold for TSV evidence is not appropriate. Instead, the ILP in SQUID sets concordant and discordant alignments into competition and selects the winner as the most reliable TSVs.

### 2.3 SQUID is able to detect non-fusion-gene TSV on two previously-studied cell lines

Fusion gene events are a strict subset of TSVs where the two breakpoints are each within a gene region and the fused sequence corresponds to the sense strand of both genes. Fusion genes thus exclude TSV events where a gene region is fused with a intergenic region or an anti-sense strand of another gene. Nevertheless, fusion genes have been implicated (likely because of available methods to detect them) in playing a role in cancer.

To probe SQUID’s ability to detect TSVs from real data, we use two cell lines, HCC1954 and HCC1395, for which previous studies have experimentally validated predicted SVs and fusion gene events. Specifically, we compile results from Bignell et al. [34], Stephens et al. [35], Galante et al. [36], Zhao et al. [37] and Robinson et al. [38] for HCC1954, and results from Stephens et al. [35] and Zhang et al. [13] for HCC1395. After removing short deletions and overlapping structural variations among different studies, we have 326 validated structural variations for HCC1954 cell line, in which 245 of them have at least one breakpoint outside a gene region, and the rest (81) have both breakpoints within gene region; we have 256 validated true structural variations for HCC1395 cell line, in which 94 have at least one breakpoint outside a gene region, while the rest (162) have both breakpoints within gene. For a predicted structural variation to be true positive, both predicted breakpoints should be within a window of 30kb of true breakpoints and the predicted orientation should agree with the true orientation. We use a relatively large window since the true breakpoints can be located within an intron or other non-transcribed region, while the observed breakpoint from RNA-seq reads will be at a nearby coding or expressed region.

We use publicly available RNA-seq data from the NIH Sequencing Read Archive (SRA; accessions: SRR2532344 and SRR925710 for HCC1954, SRR2532336 for HCC1395). Because the data are from a pool of experiments, the sample from which RNA-seq was collected may be different from those used for experimental validation. We align reads to the reference genome using STAR. We compare the result with the top fusion-gene detection tools evaluated in Liu et al. [39] and newer software not evaluated by Liu et al. [39], specifically, SOAPfuse [20], deFuse [14], FusionCatcher [16], JAFFA [15] and INTEGRATE [15].

When restricted to fusion gene events, SQUID achieves similar precision and sensitivity compared to fusion gene detection tools (Figure 3A). SQUID has the highest accuracy in the HCC1954 cell line, with very similar sensitivity as all fusion gene detection tools. For HCC1395, SQUID is in the middle of fusion gene detection methods, while INTEGRATE [13] and JAFFA [15] are the best performers on this sample.

**Figure 3:**
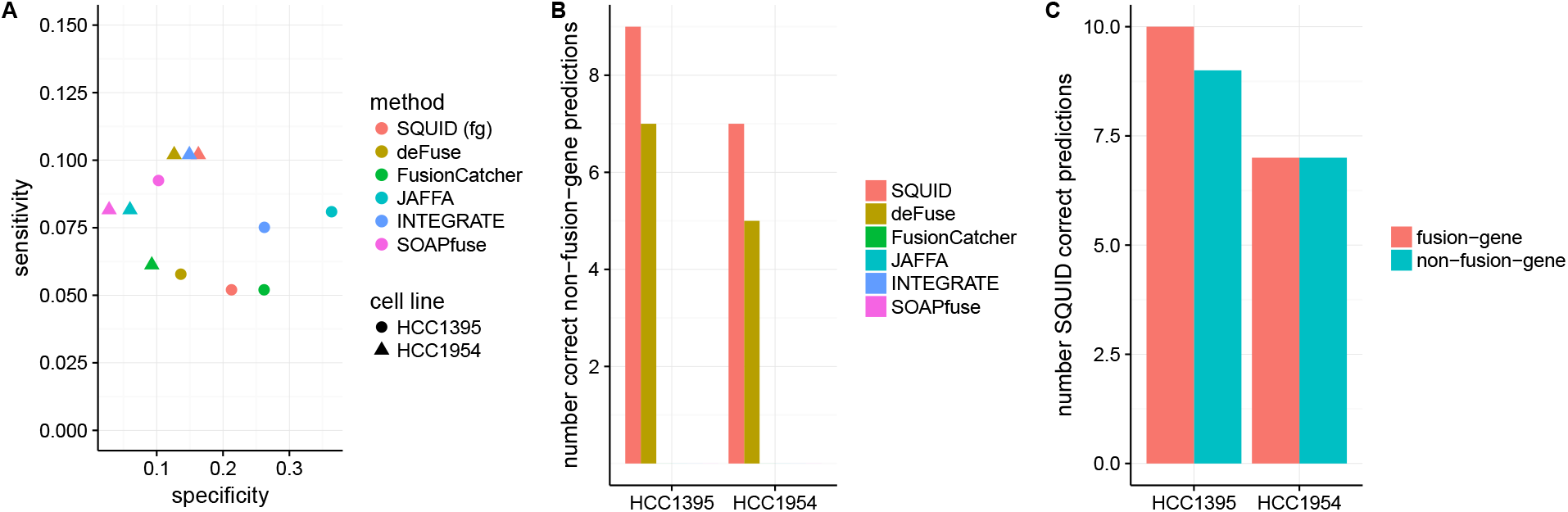
Performance of SQUID and fusion gene detection methods on breast cancer cell lines HCC1954 and HCC1395. Predictions are evaluated by previously validated SVs and fusions. (A) Sensitivity-specificity of different methods for predicting fusion gene events on both cell lines. (B) Number of correct non-fusion-gene TSV predictions that correspond to previously validated SVs. (C) Number of correctly predicted fusion-gene TSVs and non-fusion-gene TSVs from SQUID. Non-fusion-gene TSVs makes up a considerable proportion of all TSVs.

It is even harder to predict non-fusion-gene TSVs accurately, since current annotations cannot be used to limit the search space for potential read alignments or TSV events. Only SQUID and deFuse are able to detect non-fusion-gene events. Between these two methods, SQUID is able to predict more known non-fusion-gene TSVs correctly (Figure 3B). At the same time, the precision of SQUID does not decrease very much by considering both fusion-gene and non-fusion-gene TSVs (HCC1954: fusion gene specificity is 16.28%, and overall specificity is 15.56%; HCC1395: fusion gene specificity is 21.28%, and overall specificity is 19.39%). A considerable proportion of validated TSVs are non-fusion-gene TSVs: correctly predicted non-fusion-gene TSVs compose almost half of all correct predictions of SQUID (Figure 3C).

### 2.4 Charactering TSVs on four types of TCGA cancer samples

To compare the distributions and characteristics of TSVs among cancer types and between TSV types, we applied SQUID on arbitrarily selected 99 to 101 tumor samples from TCGA for each of four cancer types: breast invasive carcinoma (BRCA), bladder urothelial carcinoma (BLCA), lung adenocarcinoma (LUAD), and prostate adenocarcinoma (PRAD). (for details see Supplementary Text)

To estimate the accuracy of SQUID’s prediction on selected TCGA samples, we use WGS data of the same patients to validate TSV junctions. There are in total 72 WGS experiments available for the 400 samples (20 BLCA, 10 BRCA, 31 LUAD, 11 PRAD). For each TSV prediction, we extract a 25Kb sequence around both breakpoints and concatenate them according to the predicted TSV orientation. We then map the WGS reads against these junction sequences using SpeedSeq [33]. If a paired-end WGS read can only be mapped concordantly to a junction sequence but not the reference genome, that paired-end read is marked as supporting the TSV. If at least 3 WGS reads support a TSV, the TSV is considered as validated. Using this approach, SQUID’s overall validation rate is 88.21%, and this indicates that SQUID is quite accurate and reliable on TCGA data.

We find that most samples have ≈ 15–20 TSVs including ≈ 3–5 non-fusion-gene TSVs among all four cancer types (Figure 4A,B). BRCA has a longer tail on both sides of the distribution of TSV counts, where more samples contain a larger number of TSVs, and more samples contains a smaller number of TSVs. The same trend is observed when restricted to non-fusion-gene TSVs.

**Figure 4:**
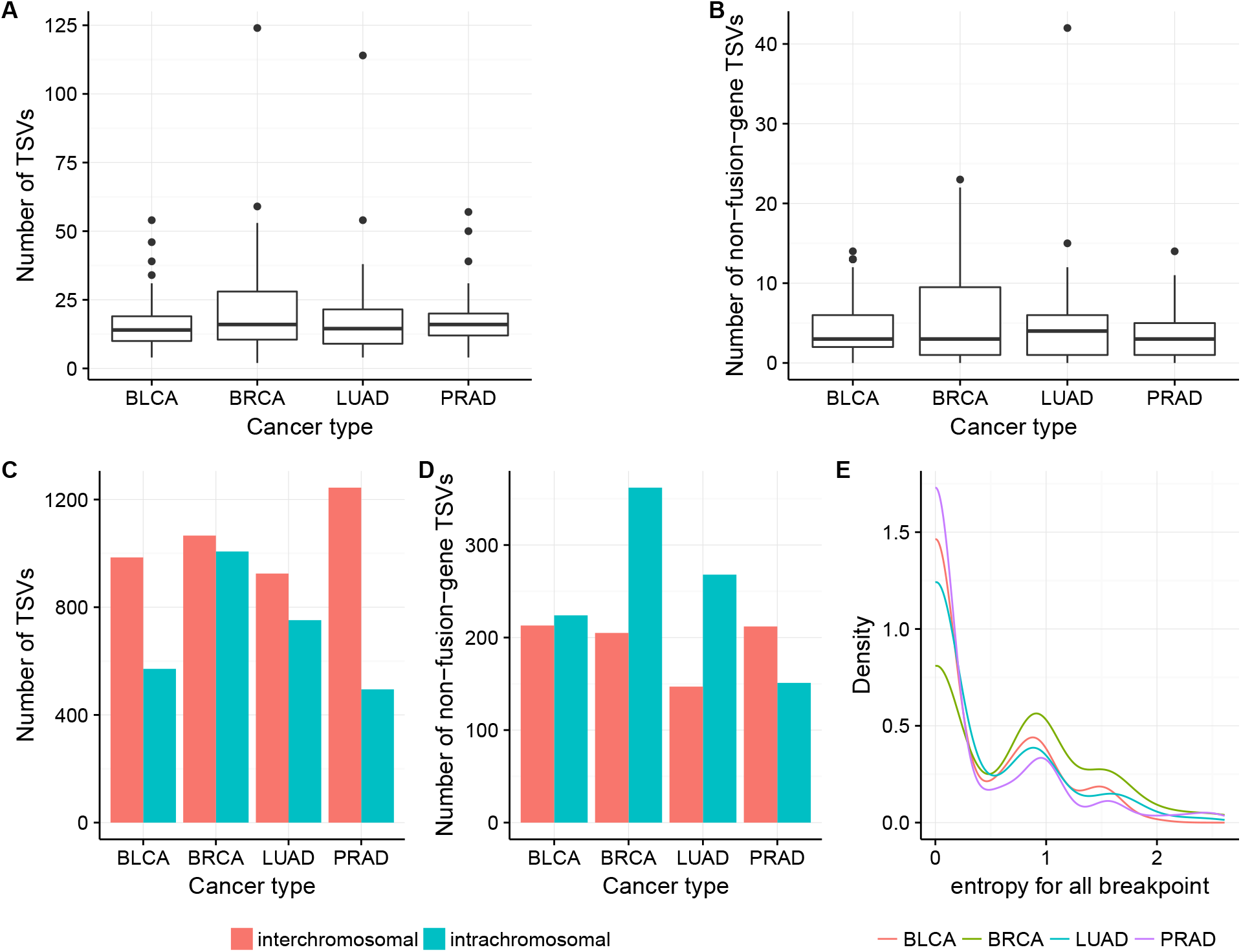
(A,B) Number of TSVs and non-fusion-gene TSVs in each sample in different cancer types. BRCA has slightly more samples with larger or smaller number of (non-fusion-gene) TSVs, thus showing a longer tail on both ends of y axis. (C,D) Number of inter-chromosomal and intra-chromosomal TSVs within all TSVs and within non-fusion-gene TSVs. Non-fusion-gene TSVs contain more intra-chromosomal events than fusion-gene TSVs. (E) For breakpoints occurring more than 3 times in the same cancer type, the distribution of the entropy of its TSV partner. The lower the entropy, the more likely the breakpoint has a fixed partner. The peak near 0 indicates a large portion of breakpoints are likely to be rejoined with the same partner in TSV. However, there are still some breakpoints that have multiple rejoined partners.

Inter-chromosomal TSVs are more prevalent than intra-chromosomal TSVs for all cancer types (Figure 4C), although this difference is much more pronounced in bladder and prostate cancer. Non-fusion-gene TSVs are more likely to have intra-chromosomal events than fusion gene TSVs (Figure 4D), and in fact in bladder, breast, and lung cancer, we detect more intra-chromosomal non-fusion-gene TSVs than inter-chromosomal non-fusion-gene TSVs. Prostate cancer is an exception in that, for non-fusion-gene TSVs, inter-chromosomal events are observed more often than intra-chromosomal events. Nevertheless, it also holds true that non-fusion-gene TSVs are more likely to be intra-chromosomal than fusion-gene, because the percentage of intra-chromosomal TSVs within non-fusion-gene TSVs is higher than that within all TSVs.

For a large proportion of breakpoints occurring multiple times within a cancer type, their partner in the TSV is likely to be fixed and to reoccur every time that breakpoint is used. To quantify this, for each breakpoint that occurred ≥ 3 times, we compute the entropy of its partner promiscuity. Specifically, we derive a discrete, empirical probability distribution of partners for each breakpoint and compute the entropy of this distribution. This measure thus represents the uncertainty of the partner given one breakpoint, with higher entropy corresponding to a less conserved partnering pattern. In Figure 4E, we see that there there is a high peak near 0 for all cancer types, which indicates that for a large proportion of recurring breakpoints, we are certain about its rejoined partner once we know the breakpoint. However, there are also promiscuous breakpoints with entropy larger than 0.5.

### 2.5 Tumor suppressor genes can undergo TSV and generate altered transcripts

Tumor suppressor genes (TSG) protect cells from becoming cancer cells. Usually their functions involve inhibiting cell cycle, facilitating apoptosis, and so on [40]. Mutations in TSGs may lead to loss of function of the corresponding proteins and benefit tumor growth. For example, homozygous loss-of-function mutation in p53 is found in about half of cancer samples across various cancer types [41]. TSVs are likely to cause loss of function of TSGs as well. Indeed, we observe several TSGs that are affected by TSVs, both of the fusion-gene type and the non-fusion-gene type.

The *ZFHX3* gene encodes a transcription factor that transactivates cyclin-dependent kinase inhibitor 1A (aka *CDKN1A*), a cell cycle inhibitor [42]. We find that in one BLCA and one BRCA sample, there are TSVs affecting *ZFHX3*. These two TSVs events are different from each other in terms of the breakpoint partner outside of *ZFHX3*. In the BLCA tumor sample, a intergenic region is inserted after the third exon of *ZFHX3* (Figure 5A). The fused transcript stops at the inserted region, causing the *ZFHX3* transcript to lose the rest of its exons. In the BRCA tumor sample, a region of the anti-sense strand of gene *MYLK3* is inserted after the third exon of *ZFHX3* gene (Figure 5B). Because codons and splicing sites are not preserved on the anti-sense strand, the transcribed insertion region does not correspond to known exons of *MYLK3* gene, but covers the range of first exon of *MYLK3* and extend to the first intron and 5’ intergenic region. Transcription stops within inserted region, and causes the *ZFHX3* transcript to lose exons after exon 3, which resembles the fusion with intergenic region in BLCA sample.

**Figure 5:**
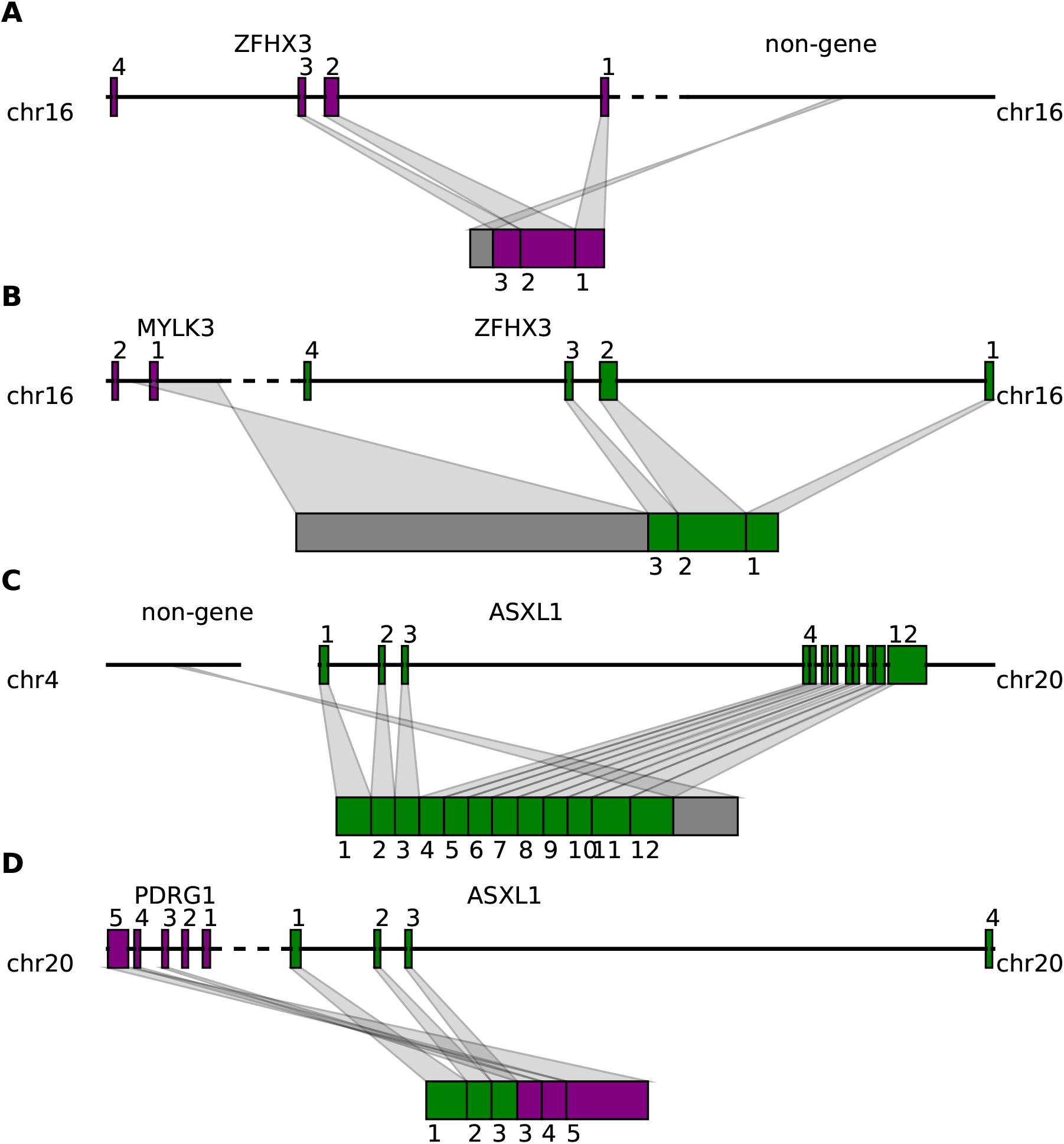
Tumor suppressor genes are affected by both fusion-gene and non-fusion-gene TSVs and generate transcripts with various features. (A) *ZFHX3* is fused with a intergenic region after exon 3. The transcript stops at the inserted region, losing the rest of exons. (B) *ZFHX3* is fused with a part of *MYLK3* anti-sense strand after exon 3. Codon and splicing signals are not preserved on anti-sense strand, thus *MYLK3* antisense insertion acts the same as intergenic region insertion, and causes transcription stop before reaching the rest of *ZFHX3* exons. (C) *ASXL1* is fused with an intergenic region in the middle of exon 12. The resulting transcript contains a truncated *ASXL1* exon 12 and intergenic sequence. (D) The first 3 exons of *ASXL1* gene are joined with last 3 exons of *PDRG1*, resulting in a fused transcript containing 6 complete exons from both *ASXL1* and *PDRG1*.

Another example is given by the *ASXL1* gene, which is essential for activating *CDKN2B* to inhibit tumor-genesis [43]. We observe two distinct TSVs related to *ASXL1* from BLCA and BRCA samples. The first TSV merges the first 11 exons and half of exon 12 of *ASXL1* with a intergenic region on chromosome 4 (Figure 5C). Transcription stops at the inserted intergenic region, leaving the rest of exon 12 not transcribed. The breakpoint within the *ASXL1* is before the 3’ UTR, so the downstream protein sequence from exon 12 will be affected. The other TSV involving *ASXL1* is a typical fusion-gene TSV where the first three exons of *ASXL1* are fused with the last three exons from the *PDRG1* gene (Figure 5D). Protein domains after *ASXL1* exon 4 and before *PDRG1* exon 2 are lost in the fused transcript.

These non-fusion-gene examples are novel predicted TSV events that are not typically detectable via traditional fusion-gene detection methods using RNA-seq data. They suggest that non-fusion-gene events can also be involved in tumorgenesis by causing disruption of tumor suppressor genes.

## 3 Discussion

We developed SQUID, the first algorithm for accurate and comprehensive TSV detection that targets both traditional fusion-gene detection and the much broader class of general TSVs. SQUID exhibits far higher precision at similar sensitivities compared with WGS-based SV detection methods and pipelines of de novo transcriptome assembly and transcript-to-genome alignment. In addition, it has the ability to detect non-fusion-gene TSVs. These features are derived from its unique approach to predicting TSVs, whereby it constructs a consistent model of the underlying rearranged genome that explains as much of the data as possible. In particular, it simultaneously considers both concordant and discordant reads, and by rearranging genome segments to maximize the number of concordant reads, SQUID generates a set of compatible TSVs that are most reliable in terms of the numbers of reads supporting them. Instead of a universal read support threshold, the objective function in SQUID naturally balances reads supporting and not supporting a candidate TSV. This design is efficient in filtering out sequencing and alignment noise in RNA-seq, especially in the annotation-free context for predicting non-fusion-gene TSV events.

We use SQUID to analyze TCGA RNA-seq data of tumor samples. We identify BRCA to have a flatter distribution of number of per-sample TSVs than the other cancer types studied. We observe that non-fusion-gene TSVs are more likely to be intra-chromosomal events than fusion-gene TSVs. This is likely due to the different sequence composition features in gene vs. non-gene regions. PRAD also stands out because the percentage of inter-chromosomal TSVs is the largest. Overall, these findings continue to suggest that different cancer types have different preferred patterns of TSVs, although the question remains whether these differences will hold up as more samples are analyzed and whether the different patterns are causal, correlated, or mostly due to non-functional randomness.

We also use SQUID to observe both non-fusion-gene and fusion-gene TSVs involving known tumor suppressor genes *ZFHX3* and *ASXL1*. In these cases, transcription usually stops within the inserted region of the non-fusion-gene TSVs, which causes the TSG transcript to lose some of its exons, reasonably leading to downstream loss of function.

Other important uses and implications for general TSVs have yet to be explored and represent possible directions for future work. TSVs will impact accuracy of transcriptome assembly and expression quantification, and methodological advancements are needed to correct those downstream analyses for the effect of TSVs. For example, current reference-based transcriptome assemblers are not able to assemble from different chromosomes to handle the case of inter-chromosomal TSVs. In addition, expression levels of TSV-affected transcripts cannot be quantified if they are not present in the transcript database. Incorporating TSVs into transcriptome assembly and expression quantification can potentially improve their accuracy. SQUID’s ability to provide a new genome sequence that is as consistent as possible with the observed reads will facilitate its use as a pre-processing step for transcriptome assembly and expression quantification, though optimizing this pipeline remains a task for future work.

Several natural directions exist for extending SQUID. First, SQUID is not able to predict small deletions, instead, it treats the small deletions the same as introns. This is to some extent a limitation of using RNA-seq data: introns and deletions are difficult to distinguish, as both result in concordant split reads or stretched mate pairs. The use of gene annotations could somewhat address this problem. Second, when the RNA-seq reads are derived from a highly heterogeneous sample, SQUID is likely not able to predict all TSVs occurring in the same region if they are conflicting since it seeks a single, consistent genome model. Instead, SQUID will only pick the dominating one that is compatible with other predicted TSVs. One approach to handle this would be to iteratively re-run SQUID, removing reads that are explained at each step. Again, this represents an attractive avenue for future work.

SQUID is open source and available at http://www.github.com/Kingsford-Group/squid and the scripts to replicate the computational experiments described here are available at http://www.github.com/Kingsford-Group/ squidtest.

## 4 Methods

### 4.1 The computational problem: rearrangement of genome segments

We formulate the TSV detection problem as the optimization problem of rearranging genome segments to maximize the number of observed reads that are consistent (termed *concordant*) with the rearranged genome. This approach requires defining the genome segments that can be independently rearranged. It also requires defining what reads are consistent with a particular arrangement of the segments. We will encode both of these (segments and read consistency) within a *Genome Segment Graph* (GSG). See Figure 6 as an example.

**Figure 6:**
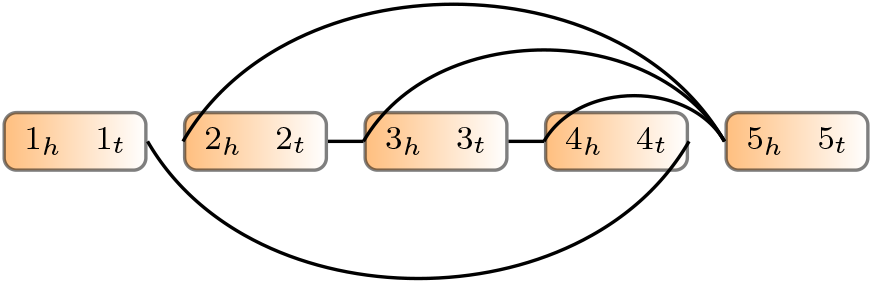
Example of genome segment graph. Boxes are genome segments, each of which has two ends subscripted by *h* and *t*. The color gradient indicates the orientation from head to tail. Edges connect ends of genome segments.

#### Definition 1 (Segment).

*A segment is a pair s* = (*S_h_, s_t_*), *where s represents a continuous sequence in reference genome and S_h_ represents its head and St represents its tail in reference genome coordinates. In practice, segments will be derived from the read locations (Section 4.4)*.

#### Definition 2 (Genome Graph (GSG)).

*A genome segment graph G* = (*V, E, w*) *is an undirected weighted graph, where V contains both endpoints of each segment in a set of segments S, i.e., V* = {*s_h_*: *s ∈ S*} ∪ {*s_t_*: *s ∈ S*}. *Thus, each vertex in the GSG represents a location in the genome. An edge* (*u,v*) *∈ E indicates that there is evidence that the location u is in fact adjacent to location v. Weight function, 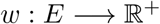, represents the reliability of an edge. Generally speaking, the weight is the number of read alignments supporting the edge, but we allow a multiplier to calculate edge weight which will be discussed below. In practice, E and w will be derived from split-aligned and paired-end reads (Section 4.5)*.

Defining vertices by endpoints of segments is required to avoid ambiguity. Only knowing that segment *i* is connected with segment *j* is not enough to recover the sequence, since different relative positions of *i* and *j* spell out different sequences. Instead, for example, an edge (*i_t_, j_h_*) indicates that the tail of segment *i* is connected head of segment *j*, and this specifies a unique desired local sequence with only another possibility of the reverse complement (i.e. it could be that the true sequence is *i · j* or *rev(j) · rev(i)*; here · indicates concatenation and *rev(i)* is the reverse complement of segment *i*).

The GSG is similar to the breakpoint graph [44] but with critical differences. A breakpoint graph has edges representing both connections in reference genome and in target genome. While edges in the GSG only represents the target genome, and they can be either concordant or discordant. In addition, the GSG does not require that the degree of every vertex is two, and thus alternative splicing and eưoneous edges can exist in the GSG.

Our goal is to reorder and reorient the segments in *S* so that as many edges in *G* are compatible with the rearranged genome as possible.

#### Definition 3 (Permutation).

*A permutation π on a set of segments S projects a segment in S to a set of integers from 1 to |S| (the size of S) representing the indices of the segments in an ordering of S. In other words, each permutation π defines a new order of segments in S*.

#### Definition 4 (Orientation Function).

*An orientation function f maps both ends of segments to 0 or 1:*

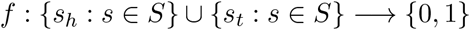

*subject to f(s_h_) + f(s_t_) = 1 for all s = (s_h_, s_t_) ∈ S. An orientation function specifies the orientations of all segments in S. Specifically, f(s_h_) = 1 means s_h_ goes first and s_t_ next, corresponding to forward strand of segment, and f(s_t_) = 1 corresponds to the reverse strand of the segment*.

With a permutation *π* and an orientation function *f*, the exact and unique sequence of genome is determined. The reference genome also corresponds to a permutation and an orientation function, where the permutation is the identity permutation, and the orientation function maps all *s_h_* to 1 and all *s_t_* to 0.

#### Definition 5 (Edge Compatibility).

*Given a set of segments S, a genome segment graph G = (V, E, w), a permutation π on S, and an orientation function f, an edge e = (u_i_,V_j_) ∈ E, where u_i_ ∈ {u_h_, u_t_} and V_j_ ∈ {v_h_, v_t_}, is compatible with permutation π and orientation f if and only if*

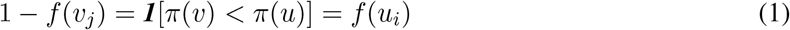

*where **1**[x] is the indicator function that is* 1 *if x is true and* 0 *otherwise. We write e ~ (π, f) if e is compatible with π and f*.

The above two edge compatibility equations (1) require that, in order for an edge to be compatible with the rearranged and reoriented sequence determined by *π* and *f*, the edge needs to connect the right side of the segment in front to the left side of segment following it. As we will see in Section 4.5, edges of GSG are derived from reads alignments. An edge being compatible with *π* and *f* is essentially equivalent to the statement that the corresponding read alignments are concordant (Section 4.3) with respect to the target genome determined by π and f. When (*π, f*) is clear, we refer to edges that are compatible as concordant edges, and edges that are incompatible as discordant edges.

With the above definitions, we formulate an optimization problem as follows:

#### Problem 1.

***Input:*** *A set of segments S and a GSG G = (V, E, w)*.

***Output:*** *Permutation π on S and orientation function f that maximizes:*

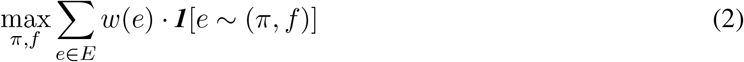

This objective function tries to find a rearrangement of genome segments (*π, f*), such that when aligning reads to the rearranged sequence, as many reads as possible will be aligned concordantly. This objective function includes both concordant alignments and discordant alignments and sets them in competition, which will be effective in reducing false positives when tumor transcripts out-number normal transcripts. There is the possibility that some rearranged tumor transcripts are out-numbered by normal counterparts. In order to be able to detect TSV in this case, depending on the setting, we may weight discordant read alignments more than concordant read alignments. Specifically, for each discordant edge *e*, we multiply the weight *w(e)* by a constant *α*, which represents our estimate of the ratio of normal transcripts over tumor counterparts.

The final TSVs are modeled as pairs of breakpoints. Denote the permutation and orientation corresponding to an optimally rearranged genome as (*π*, f\**) and those that correspond to reference genome as (*π_0_, f_0_*). An edge *e* can be predicted as a TSV if *e ~ (π*, f*)* and *e ≁ (π_0_, f_0_)*.

### 4.2 Integer linear programming formulation

We use integer linear programming (ILP) to compute an optimal solution *(π*, f*)* of Problem 1. To do this, we introduce the following boolean variables:

- *x_e_*: *x_e_* = 1 if edge *e ~ (π*, f*)*, and *x_e_* = 0 if not.
- *z_uv_*: *z_uv_* = 1 if segment *u* is before v in the permutation π*, and 0 otherwise.
- *y_u_*:*y_u_* = 1 if *f* (u_h_)* = 1 for segment *u*.

With this representation, the objective function can be rewritten as

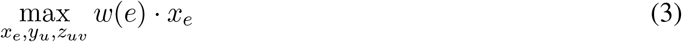

We add constraints to the ILP derived from edge compatibility equations (1). Without loss of generality, we first suppose segment *u* is in front of *v* in the reference genome, and edge *e* connects *u_t_* and *v_h_* (which is a tail-head connection). Plugging in *u_t_*, the first equation in (1) is equivalent to 1 – **1**[*π*(*u*) *> π*(*v*)] = 1 – *f*(*u_t_*), and can be rewritten as **1**[*π*(*u*) *< π*(*v*)] = *f*(*u_h_*) = *y_u_*. Note that **1**[*π*(*u*) *< π*(*v*)] has the same meaning as *z_uv_*; it leads to the constraint *z_uv_ = y_u_*. Similarly, the second equation in (1) indicates *z_uv_ = y_v_*. Therefore, *x_e_* can only reach 1 when *y_u_ = y_v_ = z_uv_*. This is equivalent to the inequalities (4) below. Analogously, we can write constraints for other three types of edge connections: tail-tail connections impose inequalities (5); head-head connections impose inequalities (6); head-tail connections impose inequalities (7):

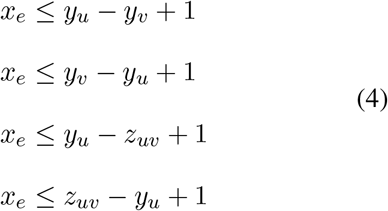

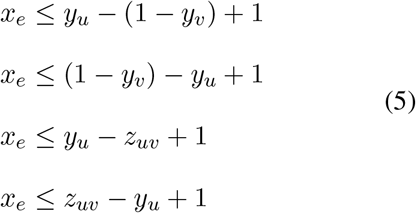

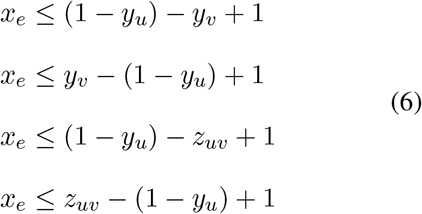

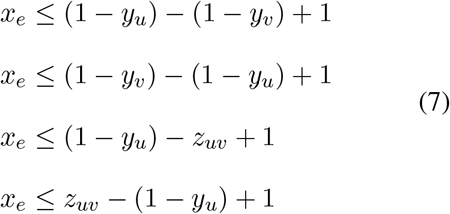

We also add constraints to enforce that *z_uv_* forms a valid topological ordering. For each pair of nodes *u* and *v*, one must be in front of other, that is *z_uv_ + z_vu_* = 1. In addition, for each triple of nodes, *u, v* and *w*, they cannot be all in front of another; one must be at the beginning of these three and one must be at the end.

Therefore we add 1 ≤ *z_uv_ + z_vw_ + z_wu_* ≤ 2.

Solving an ILP in theory takes exponential time, but in practice, solving the above ILP to rearrange genome segments is very efficient. The key is that we can solve for each connected component separately. Because the objective maximizes the sum of compatible edge weights, the best rearrangement of one connected component is independent from the rearrangement of another because by definition there are no edges between connected components.

### 4.3 Concordant and discordant alignments

Discordant alignments are alignments of reads that contradict library preparation in sequencing. Concordant alignments are alignments of reads that agree with the library preparation. Take Illumina sequencing as an example. In order for a paired-end read alignment to be concordant, one end should be aligned to the forward strand and the other to the reverse strand, and the forward strand aligning position should be in front of the reverse strand aligning position (Figure 7A). Concordant alignment traditionally used in WGS also requires that a read cannot be split and aligned to different locations. But these requirements are invalid in RNA-seq alignments because alignments of reads can be separated by an intron with unknown length.

**Figure 7:**
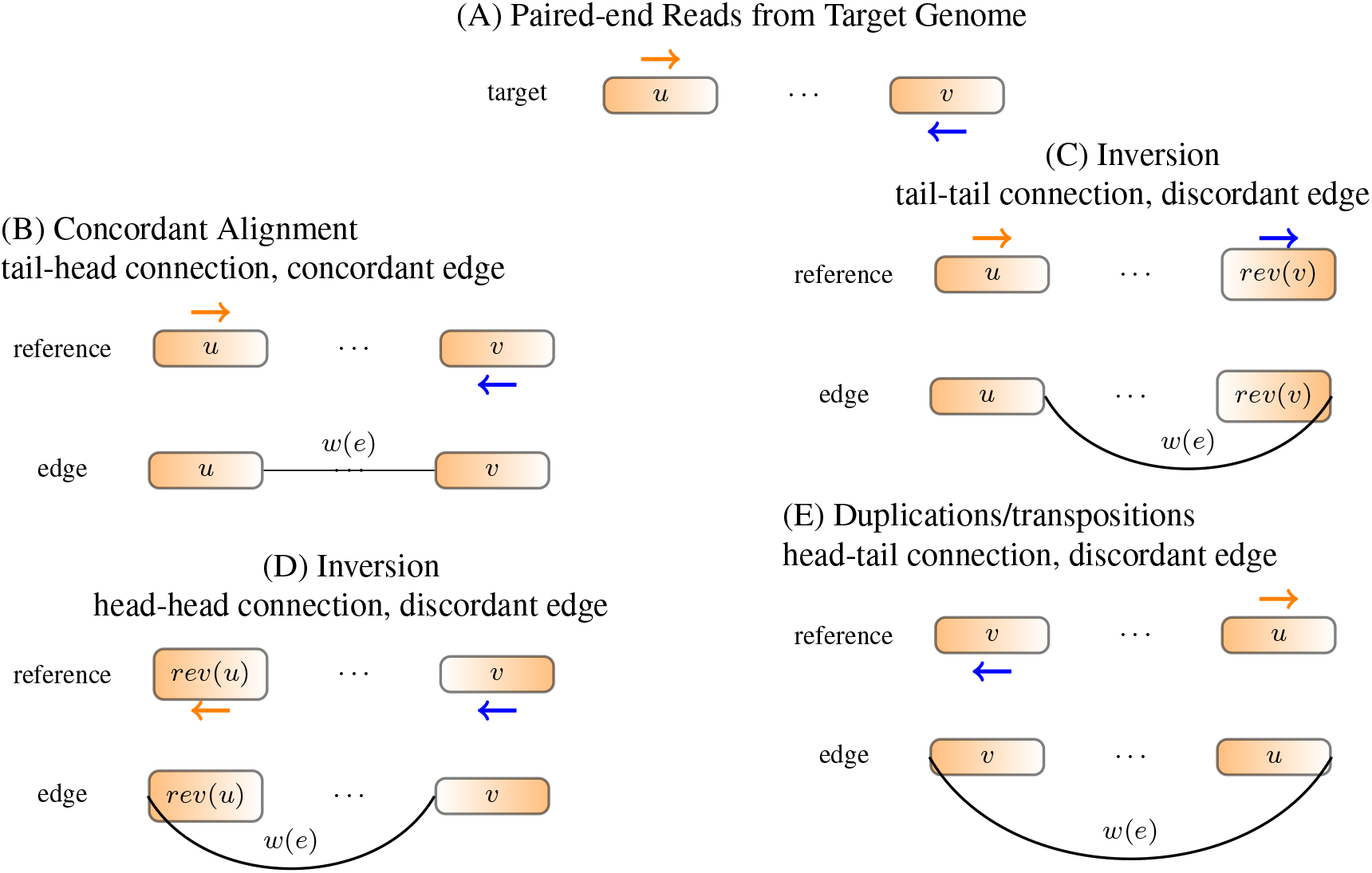
Constructing edges from alignment. (A) Read positions and orientations generated from the target genome. (B) If the reference genome does not have rearrangements, the read should be concordantly aligned to reference genome. An edge is added to connect the right end of *u* to the left end of *v*. Traversing the two segments along the edge reads out *u · v*, which is the same as reference. (C) Both ends of the read align to forward strand. An edge is added to connect the right end of *u* to the right end of *rev(v)*. Traversing the segments along the edge reads out sequence *u · rev* {*rev*(*v*)) = *u · v*, which recovers the target sequence and the read can be concordantly aligned to. (D) If both ends align to the reverse strand, an edge is added to connect the left end of front segment to the left end of back segment. (E) If two ends of a read point out of each other, an edge is added to connect the left end of front segment to the right end of back segment.

We define concordance criteria separately for split-alignment and paired-end alignment. If one end of a paired-end read is split into several parts and each part is aligned to a location, the end has split-alignments. Denote the vector of the split alignments of an end to be *R* = [*A_1_, A_2_, …, A_r_*] (*r* depends on the number of splits). Each alignment *R*[*i*] = *A_i_* is comprised of 4 components: chromosome (Chr), alignment starting position (Spos), alignment ending position and orientation (Ori, with value either + or –). We require that the alignments *A_i_* are sorted by their position in read. A split-aligned end *R* = [*A_1_, A_2_, …, A_r_*] is concordant if all the following conditions hold:

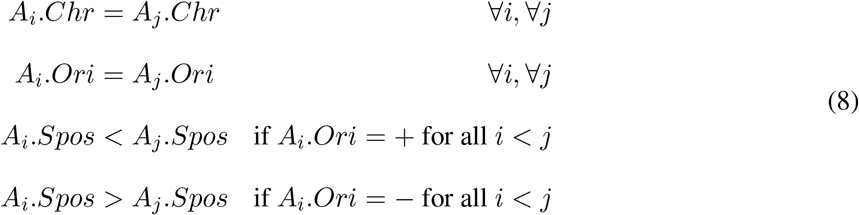

If the end is not split, but continuous aligned, the alignment automatically satisfies equation (8). Denote the alignments of *R*’s mate as *M =* [*B_1_, B_2_, …, B_m_*]. An alignment of the paired-end read is concordant if the following conditions all hold:

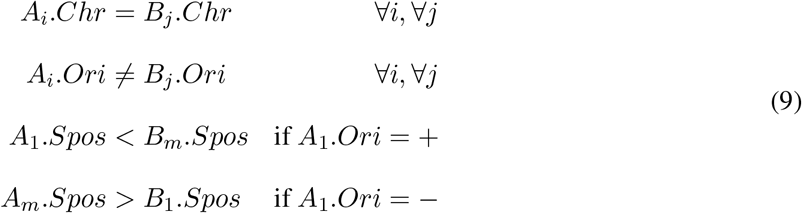

We only require the left-most split of the forward read *R* be in front of the left-most split of the reverse read *M* since the two ends in a read pair may overlap. In order for a paired-end read to be concordant, each end should satisfy split-read alignment concordance (8), and the pair should satisfy paired-end alignment concordance (9).

### 4.4 Splitting the genome into segments *S*

We use a set of breakpoints to partition the genome. The set of breakpoints contains two types of positions: (1) the start position and end position of each interval of overlapping discordant alignments, (2) an arbitrary position in each 0-coverage region.

Ideally, both ends of a discordant read should be located in separate segments, otherwise, the discordant read contained in a single segment will always be discordant no matter how the segments are rearranged. Assuming discordant read alignments of each TSV pile up around the breakpoints and do not overlap with discordant alignments of other TSVs, we set a breakpoint on the start and end positions of each contiguous interval of overlapping discordant alignments.

For each segment that contains discordant read alignments, it may also contain concordant alignments that connect the segment to its adjacent segments. To avoid having all segments in GSG connected to their adjacent segments and thus creating one big connected component, we pick the starting point of each 0-coverage region as a breakpoint. By adding those breakpoint, different genes will be in separate connected components unless some discordant reads support their connection. Overall, the size of each connected component is not very large: the number of nodes generated by each gene is approximately the number of exons located in them and these gene subgraphs are connected only when there is a potential TSV between them.

### 4.5 Defining edges in the genome segment graph

In a GSG, an edge is added between two vertices when there are reads supporting the connection. For each read spanning different segments, we build an edge such that when traversing the segments along the edge, the read is concordant with the new sequence (equations (8) and (9)). Examples of deriving an edge from a read alignment are given in Figure 7. In this way, concordance of an alignment and compatibility of an edge with respect to a genome sequence is equivalent.

The weight of a concordant edge is the number of read alignments supporting the connection, while the weight of a discordant edge is the number of supporting alignments multiplied by discordant edge weight coefficient *α*. Edges with very low read support are likely to be a result of alignment error, therefore we filter out edges with weight lower than a threshold *θ*. Segments with too many connections to other regions are likely to have low mappability, so we also filter out segments connecting to more than *γ* other segments. The parameters *α, θ*, and *γ* are the most important user-defined parameters to SQUID (Supplementary Table S1 and Supplementary Figure S2).

### 4.6 Identifying TSV breakpoint locations

Edges that are discordant in the reference genome indicate potential rearrangements in transcripts. Among those edges, some are compatible with the permutation and orientation from ILP. These edges are taken to be the predicted TSVs. For each edge that is discordant initially but compatible with the optimal rearrangement found by the ILP, we examine the discordant read alignments to determine the exact breakpoint located within related segments. Specifically, for each end of a discordant alignment, if there are 2 other read alignments that start or end in the same position and support the same edge, then the end of the discordant alignment is predicted to be the exact TSV breakpoint. Otherwise, the boundary of the corresponding segment will be output as the exact TSV breakpoint.

### 4.7 Simulation methodology

Simulations with randomly added structural variations and simulated RNA-seq reads were used to evaluate SQUID’s performance in situations with a known correct answer. RSVsim [45] was used to simulate SV on the human genome (Ensembl 87 or hg38) [46]. We use the 5 longest chromosomes for simulation (chromosome 1 to chromosome 5). RSVsim introduces 5 different types of SVs: deletion, inversion, insertion, duplication, and inter-chromosomal translocation. To vary the complexity of the resulting inference problem, we simulated genomes with 200 SVs of each type, 500 SVs of each type, and 800 SVs of each type. We generated 4 replicates for each level of SV complexity (200, 500, 800). For inter-chromosomal translocations, we only simulate 2 events because only 5 chromosomes were used.

In the simulated genome with SVs, the original gene annotations are not applicable, and we cannot simulate gene expression from the rearranged genome. Therefore, for testing purposes, we interchange the role of the reference (hg38) and rearranged genome, and use the new genome as the reference genome for alignment, and hg38 with the original annotated gene positions as the target genome for sequencing. Flux Simulator [47] was used to simulate RNA-seq reads from the hg38 genome using the Ensembl annotation version 87 [48].

After simulating SVs on genome, we need to transform the SVs into a set of TSVs, because not all SVs affect transcriptome, and thus not all SVs can be detected by RNA-seq. To derive the list of TSVs, we compare the positions of simulated SVs with the gene annotation. If a gene is affected by an SV, some adjacent nucleotides in the corresponding transcript may be located far part in the RSVsim-generated genome. The adjacent nucleotides can be consecutive nucleotides inside an exon if the breakpoint breaks the exon, or the end points of two adjacent exons if the breakpoint hits the intron. So for each SV that hits a gene, we find the pair of nucleotides that are adjacent in transcript and separated by the breakpoints, and convert them into coordinate of the RSVsim-generated genome, thus deriving the TSV.

We compare SQUID to the pipeline of de novo transcriptome assembly and transcript-to-genome alignment. We also use the same set of simulations to test whether existing WGS-based SV detection methods can be directly applied to RNA-seq data. For the de novo transcriptome assembly and transcript-to-genome alignment pipeline, we use all combinations of the existing software Trinity [23], Trans-ABySS [22], GMAP [27] and MUMmer3 [26]. For WGS-based SV detection methods, we test LUMPY [7] and DELLY2 [6]. We test both STAR [49] and SpeedSeq [33] (which is based on BWA-MEM [50]) to align RNA-seq reads to the genome. LUMPY is only compatible with SpeedSeq output, so we do not test it with STAR alignments.

## Abbreviations

ILP: integer linear programming; SV: structural variation; TSV: transcriptomic structural variation; TCGA: The Cancer Genome Atlas; WGS: whole genome sequencing.

## Acknowledgements

We thank Jacob West-Roberts for useful discussions. This research is funded in part by the Gordon and Betty Moore Foundation’s Data-Driven Discovery Initiative through Grant GBMF4554 to C.K., by the US National Science Foundation (CCF-1256087, CCF-1319998) and by the US National Institutes of Health (R21HG006913, R01HG007104), and by the Curci Foundation. C.K. received support as an Alfred P. Sloan Research Fellow. This project is funded, in part, under a grant (#4100070287) with the Pennsylvania Department of Health. The Department specifically disclaims responsibility for any analyses, interpretations or conclusions.

## Supplementary Text

All experiments here are performed with SQUID version 1.0.

### Using de novo assembly and transcript to genome alignment to predict TSV

For the pipeline of de novo transcriptome assembly and transcript-to-genome alignment, the direct output is a series of alignment pieces for each assembled transcript. To derive TSV from the pieces of alignment of each transcript, we still need to use the split-read alignment concordance criteria (8) and the edge-building approach. In the case of no TSV, equation (8) still holds, since a transcript is generated from one strand of one chromosome, without rearrangements but only deletion of introns. Any violation of(8) is treated as a TSV. Here TSVs are still able to be represented by edges in GSG, where segments are the intervals of each piece of alignment, and edges are added in the same principle that traversing segments along the edges will result in a concordant alignment of the assembled transcript. The positions of both breakpoints in a TSV are exactly the two positions linked by the discordant edge, and the orientations corresponds to the connection type of the edge.

### Processing TCGA RNA-seq data

We use STAR aligner [49] to align TCGA RNA-seq reads to Ensemble genome 87 [46] with the corresponding gene annotation. STAR aligner [49] is set with the option of outputting chimeric alignments with hanging length 15bp. The chimeric alignments generated by STAR [49] are further filtered out if the paired-end reads can be aligned concordantly by SpeedSeq aligner [33] SQUID is applied to concordant alignment generated by STAR [49] and filtered chimeric alignment. The discordant edge weight coefficient *α* is set to be 1, that is, we require tumor transcripts to dominate normal transcripts in order to predict corresponding TSVs.

A large number of fusions between immunoglobulin genes are predicted by SQUID. However, there is possibility that B cells are in the mixture of sequencing and have very high expression of immunoglobulin genes (Ig). We cannot tell whether Ig rearrangements are generated by tumor cells or B cells. Therefore, we exclude Ig TSVs during post-processing and exclude them from the descriptive statistics. Note that SQUID does not exclude Ig TSVs internally, because Ig expression and VDJ recombination have been observed to 3 exist in tumor cells, and revealing the role of Ig in tumor can deepen our understanding of cancer. When ı normal cells are removed from tumor samples, using SQUID to predict Ig TSV will help the study of Ig and 2 tumor.

### SQUID parameters

**Table S1:** Value of SQUID’s parameters used in experiments

| Symbol | Description | Value |
| --- | --- | --- |
| $\gamma$ | segment degree threshold | 4 |
| $\theta$ | edge weight threshold | 5 |
| $\alpha$ | discordant edge weight coefficient | 8 (simulation and HCC cell line), 1 (TCGA) |
| $mq$ | minimum mapping quality | 255 (STAR), 1 (SpeedSeq) |
| $pq$ | low Phred quality threshold | 4 ($p = 10^{-0.4}$) |
| $l$ | maximum allowed low Phred quality length | 10 |

Note: $mq$ , $pq$ and $l$ are controls for sequencing quality and mapping quality. If mapping quality of a read is less than threshold $mq$ , the read will not be used in edge building. If the read has a low sequencing, in terms of having more than $l$ bases of sequencing quality lower than $pq$ , the read will not be used in edge building.

**Note**: *mq, pq* and *l* are controls for sequencing quality and mapping quality. If mapping quality of a read is less then threshold *mq*, the read will not be used in edge building. If the read has a low sequencing, in terms of having more than *l* bases of sequencing quality lower than *pq*, the read will not be used in edge building.

**Figure S1:**
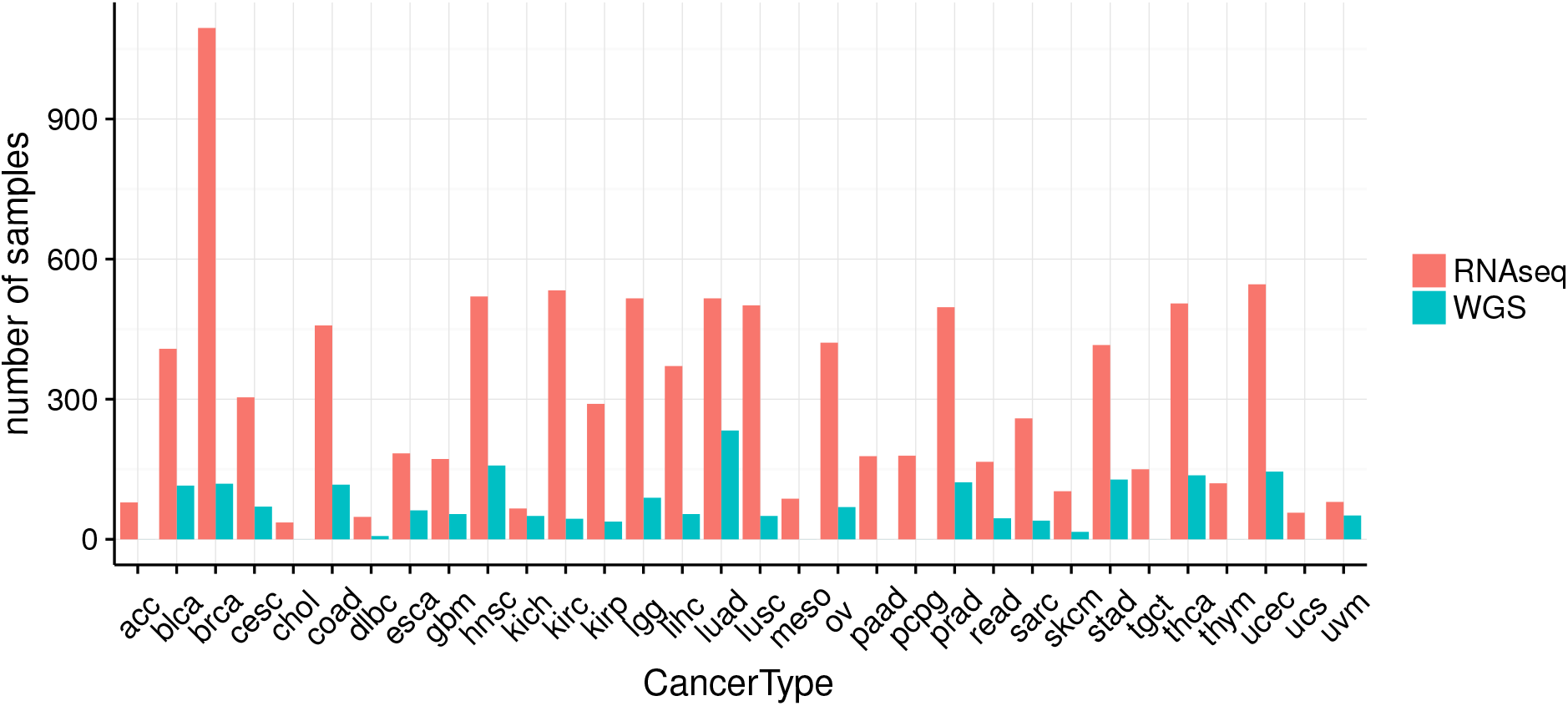
Number of samples with RNA-seq or WGS data in TCGA

**Figure S2:**
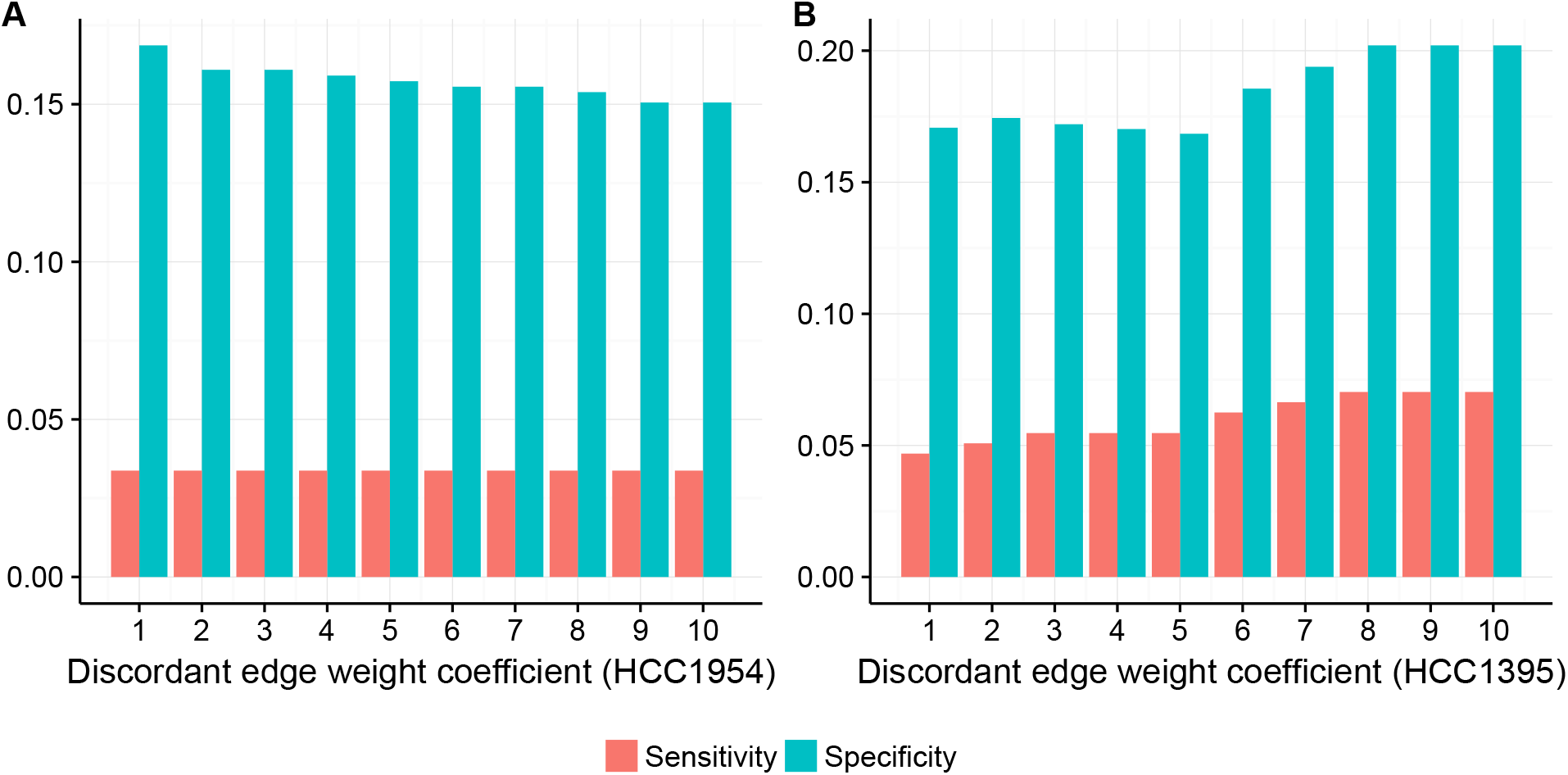
Specificity and sensitivity of SQUID against different value of discordant edge weight coefficient. (A) HCC1954 cell line. Sensitivity does not change when increasing discordant edge weight coefficient, indicating rearranged tumor transcripts out-number their normal counterparts. Specificity decreases slightly because SQUID predicts more as discordant edge weight coefficient increases. (B) HCC1395 cell line. Sensitivity and specificity reach the highest at discordant edge weight coefficient 8 and remain unchanged at 9 and 10. Some normal transcripts out-number the rearranged tumor transcripts, increasing this parameter allows SQUID to capture these TSVs.

